# How effective are contrastive learning-based approaches for activity-cliff prediction?

**DOI:** 10.64898/2026.09.08.750257

**Authors:** Akash Surendran, Ramón Alain Miranda-Quintana

## Abstract

Activity cliffs, defined as structurally similar molecules with vastly different properties represent a fundamental challenge in modern day drug discovery for property prediction models. While Graph Neural Networks (GNNs) have advanced molecular property prediction, they inherently struggle with this problem due to representation collapse and node over smoothening. In this work, we evaluate the efficacy of incorporating various contrastive learning-based loss functions, including the Supervised Contrastive (SupCon) configurations into several GNN backbones to navigate this structure-activity landscape. Tested across the MoleculeACE benchmarks, the results indicate that while contrastive losses prevent GNN mode collapse to some extent and drastically improve the property-based separation in the embedding space, their impact on absolute predictive accuracy remains highly dataset dependent. Ultimately, these methods offer critical gains in interpretability for mapping optimized lead generations, opening up new directions for models that respect these sharp pharmacological discontinuities.

## Introduction

Activity cliffs represent molecular pairs with highly similar topological or structural features exhibiting exponentially divergent biological activities against a specific therapeutic target[1–3]. In practice, these drastic shifts may be as minute as a single atom substitution or a subtle change in the hybridization state of a scaffold atom. Even such minor modifications may induce a tenfold (10x), hundredfold (100x) or even a thousandfold (1000x) shift in binding affinity against a given target, directly challenging the foundational assumptions of the molecular similarity principle. Molecular similarity principle is the central heuristic governing traditional medicinal chemistry and algorithm design, which postulates that structurally similar molecules will naturally possess comparable biological, physicochemical, and pharmacokinetic properties[4– 6]. This foundational principle drives the predictive power of Quantitative Structure-Activity Relationship (QSAR) modeling, allowing researchers to traverse the vast expanse of chemical space by interpolating biological activities of novel compounds based on their similarity to known active molecules. However, this structure-activity landscape is rarely a smooth, continuous mathematical manifold as it is frequently punctuated by severe, highly localized discontinuities or activity cliffs.

The existence of activity cliffs introduces a fascinating duality in the context of drug discovery [2], acting simultaneously as a continuous hindrance for predictive modeling while also being a pharmacological goldmine. From the perspective of classical machine learning and continuous regression algorithms, ACs manifest as troublesome outliers that violate the smoothness assumptions inherent to mathematical interpolation, often leading to predictive failures and erroneous property mapping when models attempt to smooth over these sharp anomalies[7]. Hence, in QSAR studies, these compounds are usually discarded or actively filtered out of training sets to reduce model variance. Conversely, from the perspective of a medicinal chemist, these compounds encode extraordinarily high information content as they reveal the precise limits of chemical modification, indicating exactly where a ligand clashes with an amino acid residue or disrupts a crucial hydrogen bonding network[1, 8, 9]. Hence, their identification is crucial for guiding highly optimized lead generation and avoiding late-stage failures in the drug development pipeline.

With the deployment of artificial intelligence, particularly deep learning via Graph Neural Networks (GNNs)[10– 12] became increasingly ubiquitous in molecular property prediction, the scientific community has recognized a glaring deficiency. The accurate prediction of these discontinuities remains a formidable challenge for Machine Learning and Deep Learning models as they consistently fail in this task due to representation collapse and layer-wise node over-smoothening[13, 14]. As message-passing networks aggregate neighborhood features to capture global structure, these cliff-inducing atomic perturbations get diluted into the broader shared scaffold. Furthermore, these deep networks are inherently able to learn smooth, low-frequency functions, leaving them unsuited to map the sharp discontinuities that characterize activity cliffs. This representational failure is significantly compounded by data scarcity and class imbalances, which makes these models “see” such instances as statistical outliers to avoid inflating the overall variance.

Establishing rigorous AC-specific benchmarks has been instrumental in exposing model limitations and directing methodological innovation[15]. The MoleculeACE benchmark[13] of Van Tilborg et al. consolidated 30 scaffold-stratified ChEMBL bioactivity datasets with SALI-derived[16] cliff labels, providing the first systematic framework for quantitatively evaluating cliff vs non-cliff prediction performance. The results of this benchmarking study revealed near-universal model degradation on cliff subsets across diverse target families and representations. On the other hand, this has led to scenarios where simple traditional machine learning baselines utilizing Extended Connectivity Fingerprints (ECFPs)[17] completely outcompete deep learning architectures in cliff prediction tasks.

In response to these challenging benchmarks, novel deep learning frameworks have recently emerged, leveraging specialized architectures or highly customized loss functions. For instance, to overcome the spectral bias[18] that cause standard GNNs to oversmooth the subtle structural variations of ACs, models like MapCliff-WMGR framework introduces an Independent Feature Mapping (IFM) module[19]. By utilizing sinusoidal transformations to project atomic features into a frequency rich domain and subsequently processing them through a modified graph attention network (mGraphSNNGAT), MapCliff-WMGR explicitly force the model to learn the sharp, high-frequency potency shifts. Concurrently, to address the extreme class imbalance and data scarcity when it comes to activity cliffs in a standard chemical dataset, Semi-Mol architecture[20] proposes a novel semi-supervised learning (SSL) approach. It uses an instructor model to generate pseudo labels across vast unannotated chemical spaces, coupling with a self-adaptive curriculum learning algorithm to introduce more challenging cliffs at a controlled pace, effectively preventing the GNN from overfitting the majority class of smooth and continuous SAR. Conversely, other approaches like ACANet[21] optimize the training objective itself. ACANet utilizes a novel Activity-Cliff-Awareness (ACA) triplet contrastive loss function. By dynamically mining activity cliff triplets, ACANet forces the latent space to simultaneously pull together molecules with similar activities while aggressively pushing apart the embeddings of structurally similar cliff pairs, resulting in superior performance across classification and regression tasks. Building on this, in this work, we compare the performance of contrastive learning based loss functions trained on four GNN backbones in property prediction for activity cliff molecules. We also investigate the interpretability of latent space embeddings by their ability to distinguish between similar molecules with significantly different activity to see if contrastive loss based models are able to prevent mode collapse problem of GNNs when trained on activity cliffs molecules as graphs.

## Methods

### GNN backbone and loss functions

Since the main objective of this work was to compare the effect of contrastive learning based loss functions on aiding a GNN in navigating the SAR landscape of activity cliffs, we employed a GNN backbone of Principal Neighborhood Aggregation (PNA), which is also the default backbone in ACANet[21], primarily due to better performances in comparison to other variants like Graph Convolutional Network (GCN),Graph Isomorphism Network (GIN) and Graph Attention Network (GAT). The inputs to the network are small molecule graph representations which are featurized using 39 node and 10 edge features trained on a range of loss functions. More details on the ACANet architecture can be found on the article by Zitnik et al[21] and the loss functions are discussed in detail in **Appendix** . We compared the following loss functions:

#### 1. ACANet Triplet Loss

The overall ACA loss ℒ_*ACA*_ combines the standard regression loss, typically mean absolute error (MAE) with a Triplet Soft Margin (TSM) loss term:

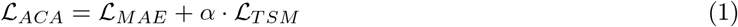

where *α* is a tunable cliff component that controls the strength of the triplet contrastive learning term ℒ_*TSM*_ . For a given batch size *N*, the first term is calculated as the MAE between the true labels *y*_*i*_ and the predicted labels *ŷ*_*i*_:

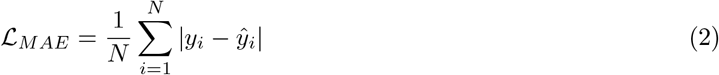

To calculate the contrastive term, the framework dynamically mines High-Value Activity Cliff Triplets (HV-ACTs) online within each batch consisting of an anchor compound(*A*), positive compound (*P*) and a negative compound. The assignment of positive or negative compounds relative to the anchor is bounded by two hyperparameters cliff lower (*c*_*l*_) and cliff upper (*c*_*u*_). Given an anchor, positives are defined by having activity difference less than lower threshold (|*y*_*A*_ − *y*_*P*_ | *< c*_*l*_) and negatives exhibit activity difference greater than the upper threshold (|*y*_*A*_ − *y*_*N*_ | *< c*_*u*_).

Once the triplets are mined, the network maps them to latent vectors (*f*) and the TSM loss penalizes the model if the latent distance between the anchor and the negative (cliff pair) is not sufficiently larger than the latent distance between the anchor and the positive (similar activity pair). For M mined high value triplets, TSM is calculated as:

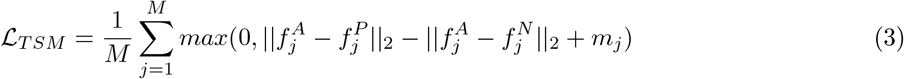

where *f* ^*A*^, *f* ^*P*^ and *f* ^*N*^ are the latent space embeddings of the anchor, positive and negative respectively and *m*_*j*_ is the soft margin applied to the triplet to accomodate a degree of tolerance based on the relative properties of the specific triplets. By pushing 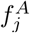 and 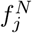 apart in latent space while maintaining proximity between 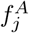 and 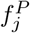, the contrastive term ensures that the GNN learns a representation space that explicitly respects the sharp discontinuities inherent to ACs.

#### 2. Propcluster SupCon Loss

We propose the following modification to the original ACA loss function, by introducing a supervised contrastive loss (SupCon)[22] term instead of the triplet soft margin loss:

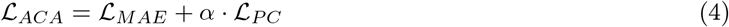

To calculate this loss term, the first step is to cluster the training set based on property value, such that for each cluster *c*_*j*_, the maximum difference in property values (in log scale) should be less than unity, i.e. 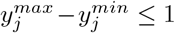. A new cluster is created and the incoming molecule is merged into it, if this condition is not satisfied. Each molecule is assigned a cluster ID, ensuring that molecules with similar activity share similar cluster and molecules with very different activities belong to different clusters. Now, *M* unique molecules are drawn at random from the full training set (across all clusters) producing a pool of size *M* (batch size). Within the pool, for every molecule *a* that can serve as an anchor:

- **Positives:** randomly draw *n*_*pos*_ ∈ [*min*_*pos*_, *max*_*pos*_] molecules from the same property cluster as *a* (excluding *a*).
- **Negatives:** randomly draw *n*_*neg*_ ∈ [*min*_*neg*_, *max*_*neg*_] molecules across different property clusters.

where we defined *min*_*pos*_*/min*_*neg*_ = 2 and *max*_*pos*_*/max*_*neg*_ = 5, allowing the model to mine for multiple instances from the positive and negative class for a given anchor (an anchor appears at most once per one sampled pool). Once these training examples are constructed, the cosine similarities between the normalized latent embeddings are calculated:

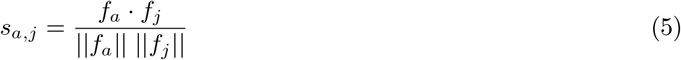

which is used to calculate the per anchor SupCon loss term, *l*_*a*_:

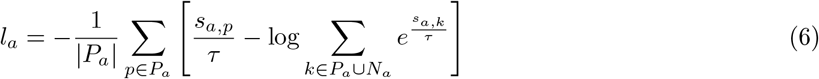

where *P*_*a*_ and *N*_*a*_ are the positive and negative class respectively for anchor *a*. The batch SupCon or the propcluster loss is calculated as the average across all valid anchors per batch:

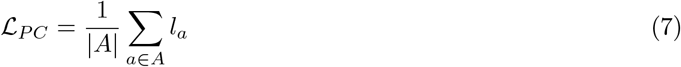

To mine harder positives/negatives, one can also implement a Tanimoto based filtering on these training examples by implementing an additional constraint for defining positives and negatives as:

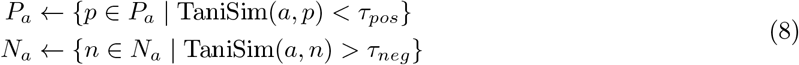

This enforces that positives are structurally dissimilar (with similar properties) and negatives are structurally similar (with very different properties) helping to mine harder training examples. The efficient training mining makes the batch processing *O*(*M*) in memory and at most *O*(*M* ^2^) in time, making it much more efficient method compared to the triplet mining.

#### 3. Pentuplet SupCon Loss

Building on the triplet loss function, we extend it to make larger training examples containing two positives and negative each per anchor:

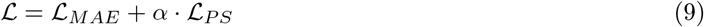

First a batch of size M is sampled from the training set, the set of positives and negatives are constructed using the property conditions (with option to implement Tanimoto based filtering) already discussed above. Valid anchors or molecules containing atleast two positives and two negatives are filtered. Let *V* be the set of valid anchors, meaning indices *i* where ∑_*j*_ *P*_*i,j*_ ≥ 2 and ∑_*j*_ *N*_*i,j*_ ≥ 2. For each anchor *i* ∈ *V*, we draw exactly 2 positives and negatives each from the set *P*_*i*_ and *N*_*i*_.

The normalized latent embeddings are then used to calculate the supervised contrastive loss (SupCon) using equation (7).

#### 4. Pentuplet Hinge Loss

Using the same procedure as the Pentuplet SupCon loss to mine the pentuples, we define the following loss function:

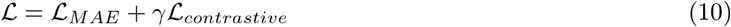

The contrastive part of the loss is composed of three terms: the cohesion, spread and anchor. The cohesion term is defined as

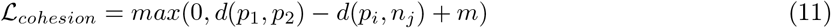

it encourages positive molecules to remain close together and farther from negatives. The spread term is defined as:

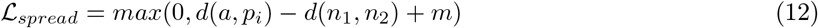

this term pushes the negative molecules to be separated from each other and from anchor-positive group. Then the anchor loss can be calculated in a similar way as the ACA triplet loss:

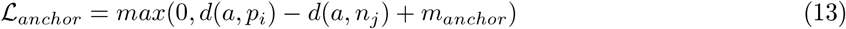

As the triplet loss, it enforces the anchor to be closer to the positives than to the negatives. The combined total contrastive loss is:

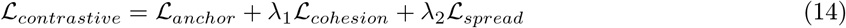

where *λ*_1_ and *λ*_2_ control the weight of the cohesion and spread terms. The contrastive loss is weighed in the final loss using the tuneable parameter *γ* like *α* for the other losses discusses above.

#### 5. Heptuplet SupCon Loss

Following the same procedure as the Pentuplet SupCon, but this time with 3 positives and negatives per anchor.

### Activity cliff benchmarking dataset and model hyperparameters

We used the 30 datasets from the MoleculeACE platform, which is a standard benchmarking database for activity cliff prediction models. Each dataset contains the binding affinity of a target protein against a set of ligands sourced from the ChEMBL 29 database. In total the dataset contains 35,632 unique molecules.

For the first step, each dataset was split into train-test splits, with batches of 128 made during each training epoch. This was followed by hyperparameter tuning specific to each method:

1. **ACANet Triplet:** cliff_lower: [0.5, 1.0, 1.5, 2.0], cliff_upper: [0.5, 1.0, 1.5, 2.0], alpha: [0.0, 0.01, 0.05, 0.1, 0.2, 0.5, 1.0], Positive similarity threshold: [0.1, 0.2, 0.3, 0.4], Negative similarity threshold: [0.6, 0.7, 0.8, 0.9]. Similarity thresholds were only used for the case when the similarity gates were activated. Each tuple contains one positive and one negative.
2. **Pentuplet SupCon:** cliff_lower: [0.5, 1.0, 1.5, 2.0], cliff_upper: [0.5, 1.0, 1.5, 2.0], alpha: [0.0, 0.01, 0.05, 0.1, 0.2, 0.5, 1.0], Temperature: [0.05, 0.07, 0.1, 0.2, 0.5], Positive similarity threshold: [0.1, 0.2, 0.3, 0.4], Negative similarity threshold: [0.6, 0.7, 0.8, 0.9]. Fixed tuple structure with 2 positives and 2 negatives.
3. **Pentuplet Hinge:** cliff_lower: [0.5, 1.0, 1.5, 2.0], cliff_upper: [0.5, 1.0, 1.5, 2.0], gamma: [0.0, 0.01, 0.05, 0.1, 0.2, 0.5, 1.0], lambda1: [0.5, 1.0, 2.0], lambda2: [0.0, 0.25, 0.5, 1.0], Positive similarity threshold: [0.1, 0.2, 0.3, 0.4], Negative similarity threshold: [0.6, 0.7, 0.8, 0.9].
4. **Heptuplet SupCon:** cliff_lower: [0.5, 1.0, 1.5, 2.0], cliff_upper: [0.5, 1.0, 1.5, 2.0], alpha: [0.0, 0.01, 0.05, 0.1, 0.2, 0.5, 1.0], Temperature: [0.05, 0.07, 0.1, 0.2, 0.5], Positive similarity threshold: [0.1, 0.2, 0.3, 0.4], Negative similarity threshold: [0.6, 0.7, 0.8, 0.9]. Same grids as the Pentuplet SupCon loss but with 3 positives and 3 negatives.
5. **Propcluster SupCon:** Clustered based on property difference of unity, so cliff_lower and cliff_upper are not needed. Alpha: [0.0, 0.01, 0.05, 0.1, 0.2, 0.5, 1.0], Temperature: [0.05, 0.07, 0.1, 0.2, 0.5], Positive similarity threshold: [0.1, 0.2, 0.3, 0.4], Negative similarity threshold: [0.6, 0.7, 0.8, 0.9]. Uses cluster aware batch sampling.

The hyperparameters are tuned in sequential fashion starting from cliff_upper/lower (not in PropCluster SupCon), Temperature or contrastive weight, alpha or hinge weights, similarity gate thresholds.

## Results

### Comparison of losses

We evaluated the prediction performance of five different loss functions: ACA Triplet Loss, Pentuplet SupCon Loss, Pentuplet Hinge Loss, Heptuplet SupCon Loss, and Propcluster SupCon Loss on 30 ChEMBL subsets with labeled binding affinities, designed to benchmark activity cliff prediction tasks. The RMSE values are reported for the overall test set as well as the cliff test set in Figure 1 for the PNA architecture. As expected, the cliff RMSE is significantly higher than the overall test set RMSE due to a higher concentration of activity cliffs, which inevitably reduces the predictive power of the model. The triplet and MAE losses show similar performance on the overall test set compared to other contrastive approaches, but the trends completely flip on the cliff test set. The losses with supervised contrastive (SupCon) component, specifically Heptuplet (0.762) and PropertyCluster (0.766) perform slightly better on the cliff subsets, proving that specialized contrastive loss functions are necessary to mitigate error on sharp structural discontinuities. The per-dataset overall test RMSE and cliff test RMSE across these loss functions are visualized in Figure 2 and 3. Clearly, the predictive performance fluctuates drastically underscoring that no single loss function universally dominates every chemical space. While the overall RMSE remains tightly grouped across the methods, cliff RMSE highlights that SupCon-based losses slightly edge out MAE when isolating the most challenging molecular pairs.

**Figure 1:**
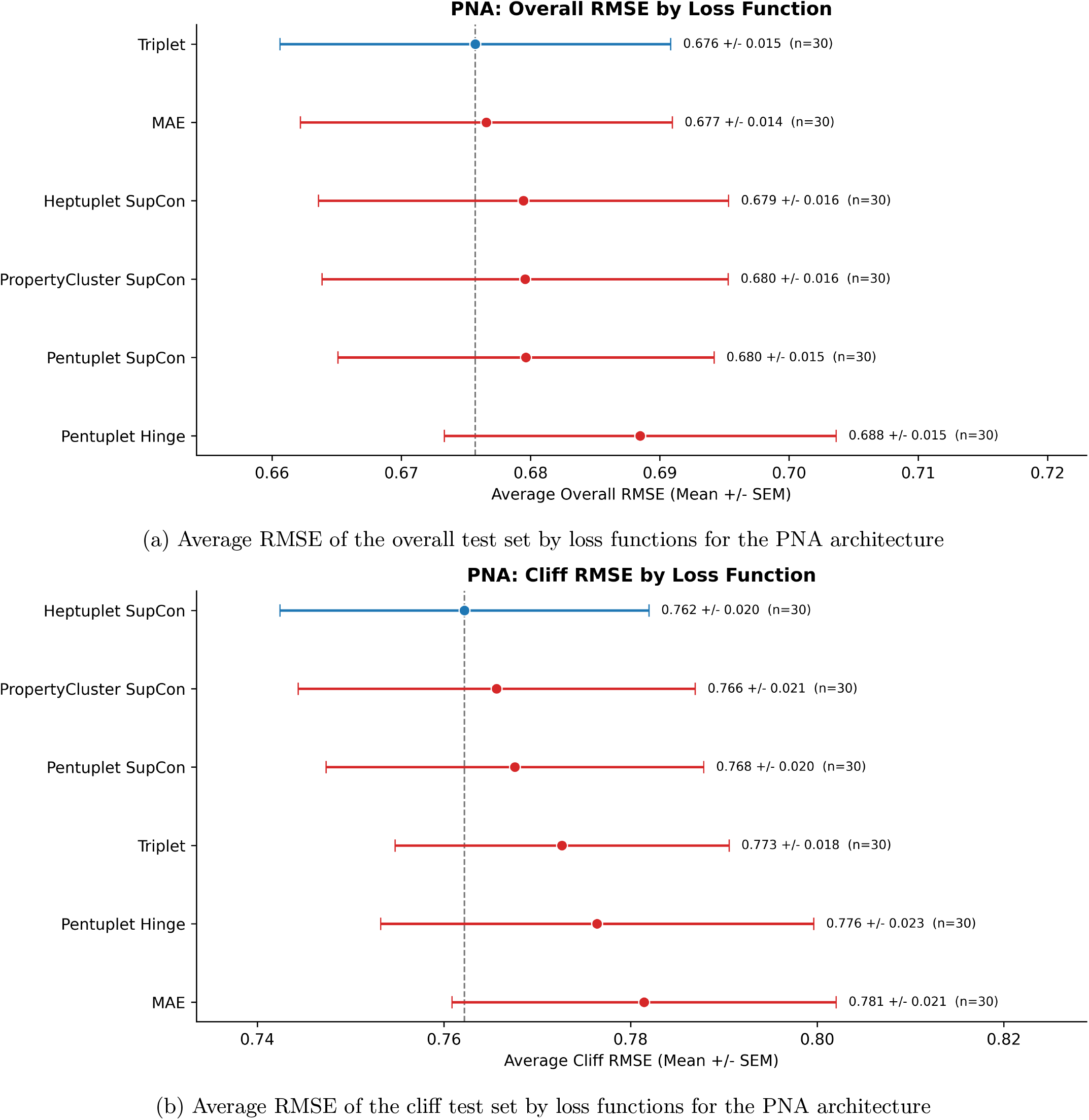
Comparison of contrastive-learning-based loss functions by overall test-set RMSE and cliff test-set RMSE across the 30 ChEMBL activity cliff datasets.

**Figure 2:**
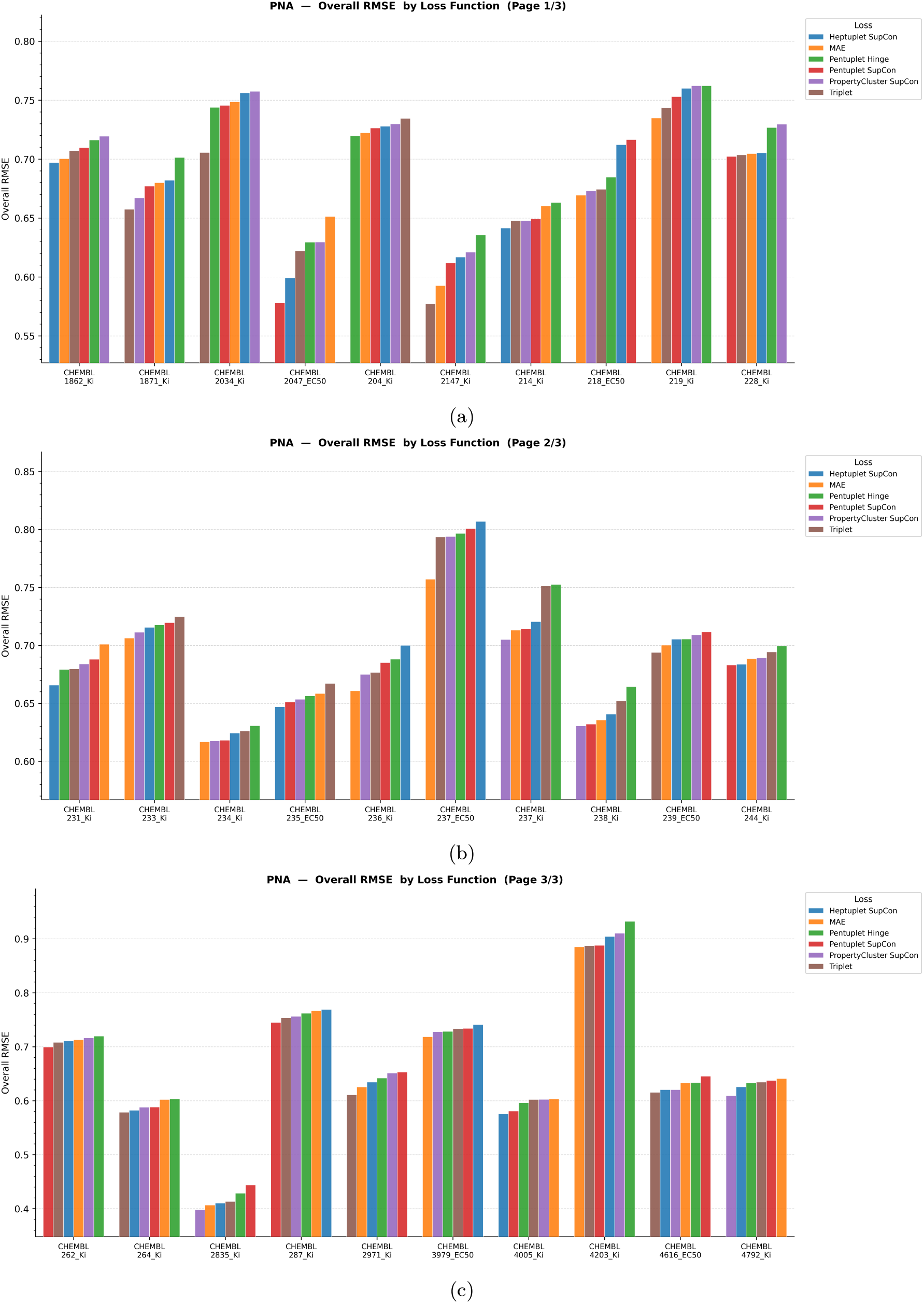
Overall test RMSE for each of the 30 ChEMBL activity cliff datasets across the PNA backbone trained on contrastive-learning based loss functions.

**Figure 3:**
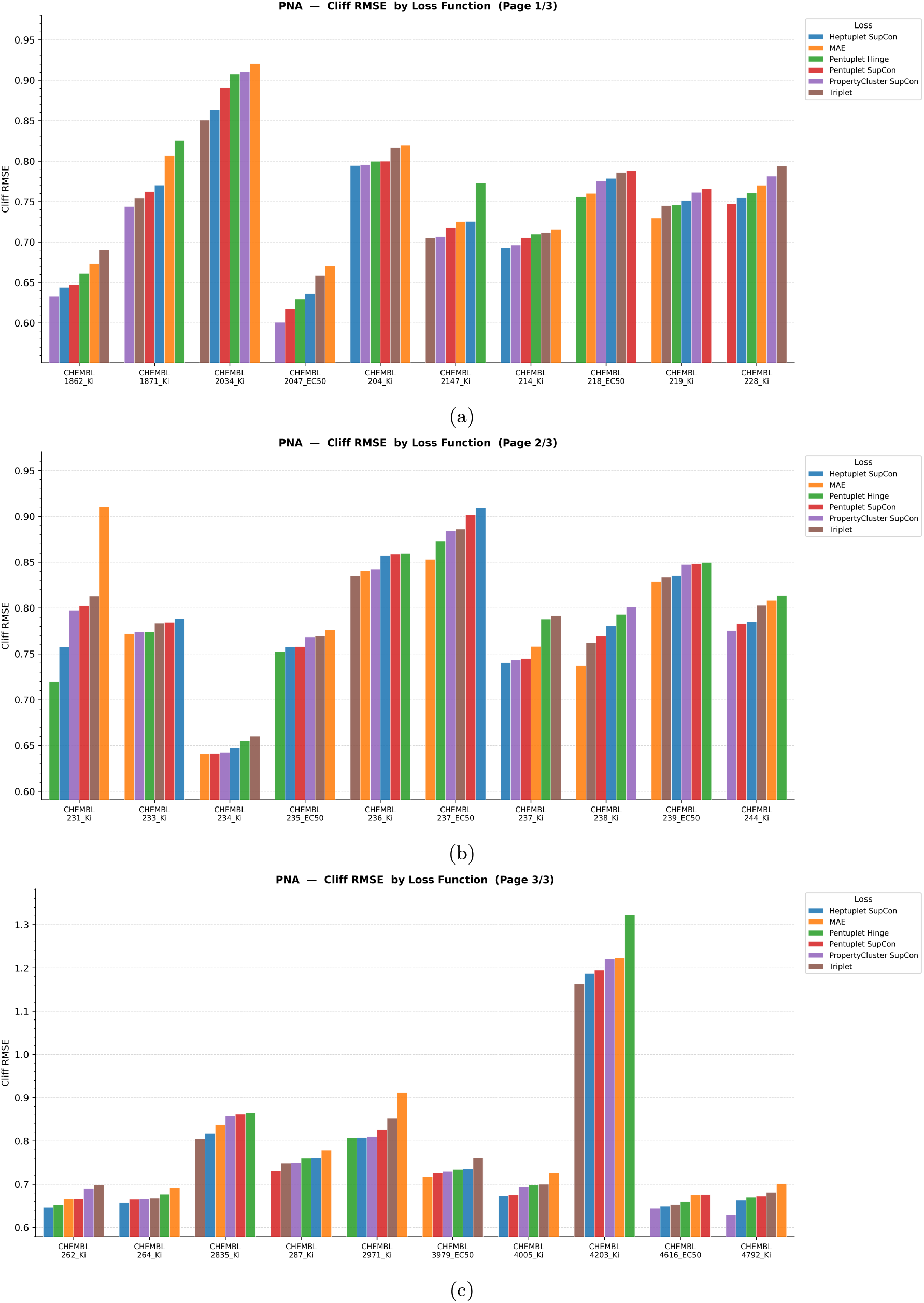
Cliff test RMSE for each of the 30 ChEMBL activity cliff datasets across the PNA backbone trained on contrastive-learning based loss functions.

### Comparisons of architectures for different losses

Now, we performed a comparison of the architecture choice for a given loss function to investigate if there is a suitable architecture for predicting activity cliffs that is consistent across all loss types. Figure 4 and 5 demonstrate the interplay between four distinct GNN architectures: PNN, GCN, GAT and GIN across the six evaluated loss functions. The overall test RMSE (Figure 4) demonstrates that the GIN architecture consistently yields the highest error margins and greatest variance across the majority of contrastive configurations, notably reaching an RMSE of 0.751 for PropertyCluster SupCon loss. In contrast, PNA and GCN backbones are the more robust structural foundations, maintaining low error baselines of 0.677 and 0.674 respectively under the MAE loss. Figure 5 mirrors these trends within the more rigorous cliff-specific test set, where absolute error rates rise naturally but the relative stability of PNA and GCN remain intact. Ultimately, this study proves that while contrastive loss functions are useful to map sharp pharmacological discontinuities, their predictive success is heavily reliant on choosing a stable architectural backbone like PNA or GCN, whereas ones like GIN prove more sensitive to the training objective.

**Figure 4:**
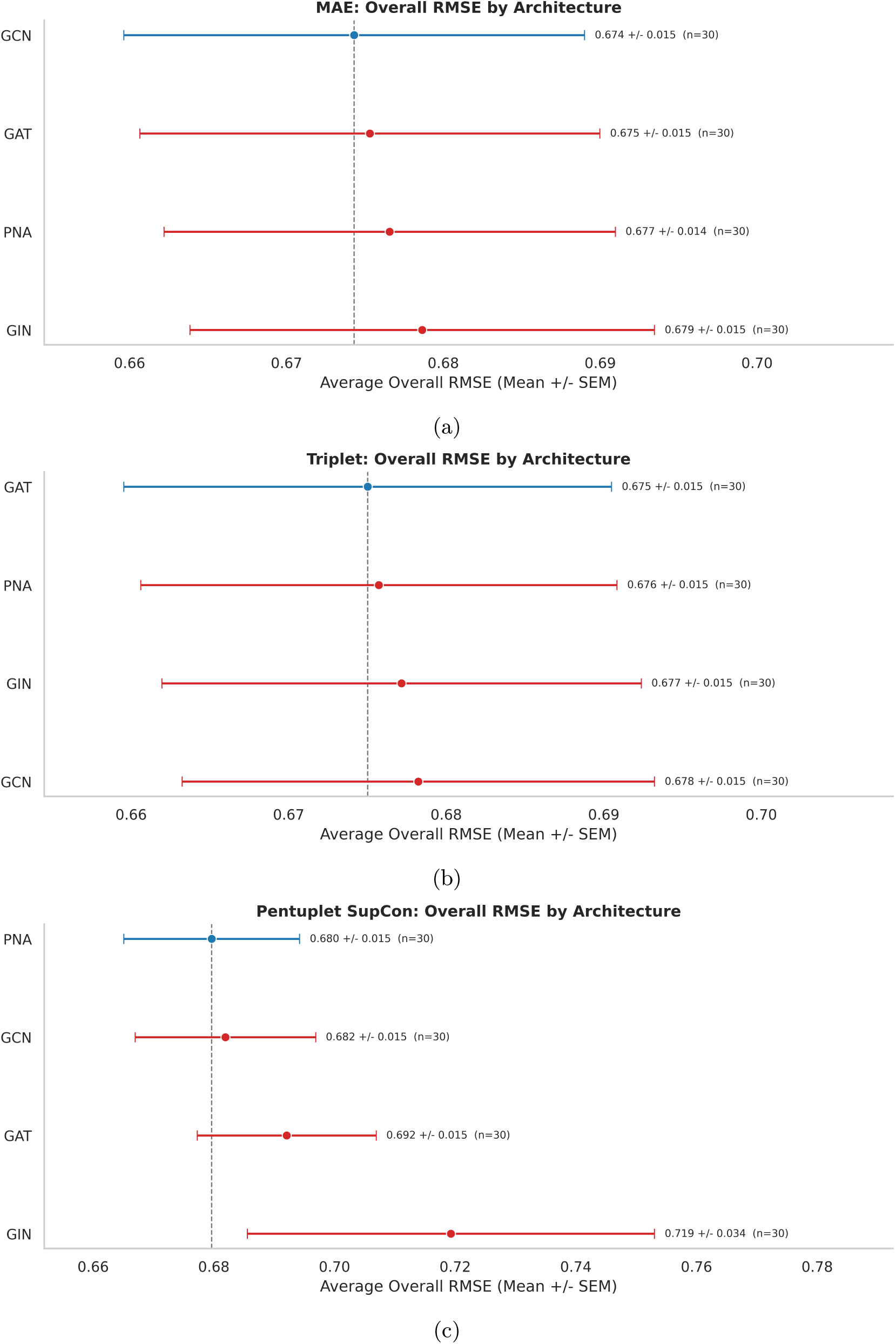
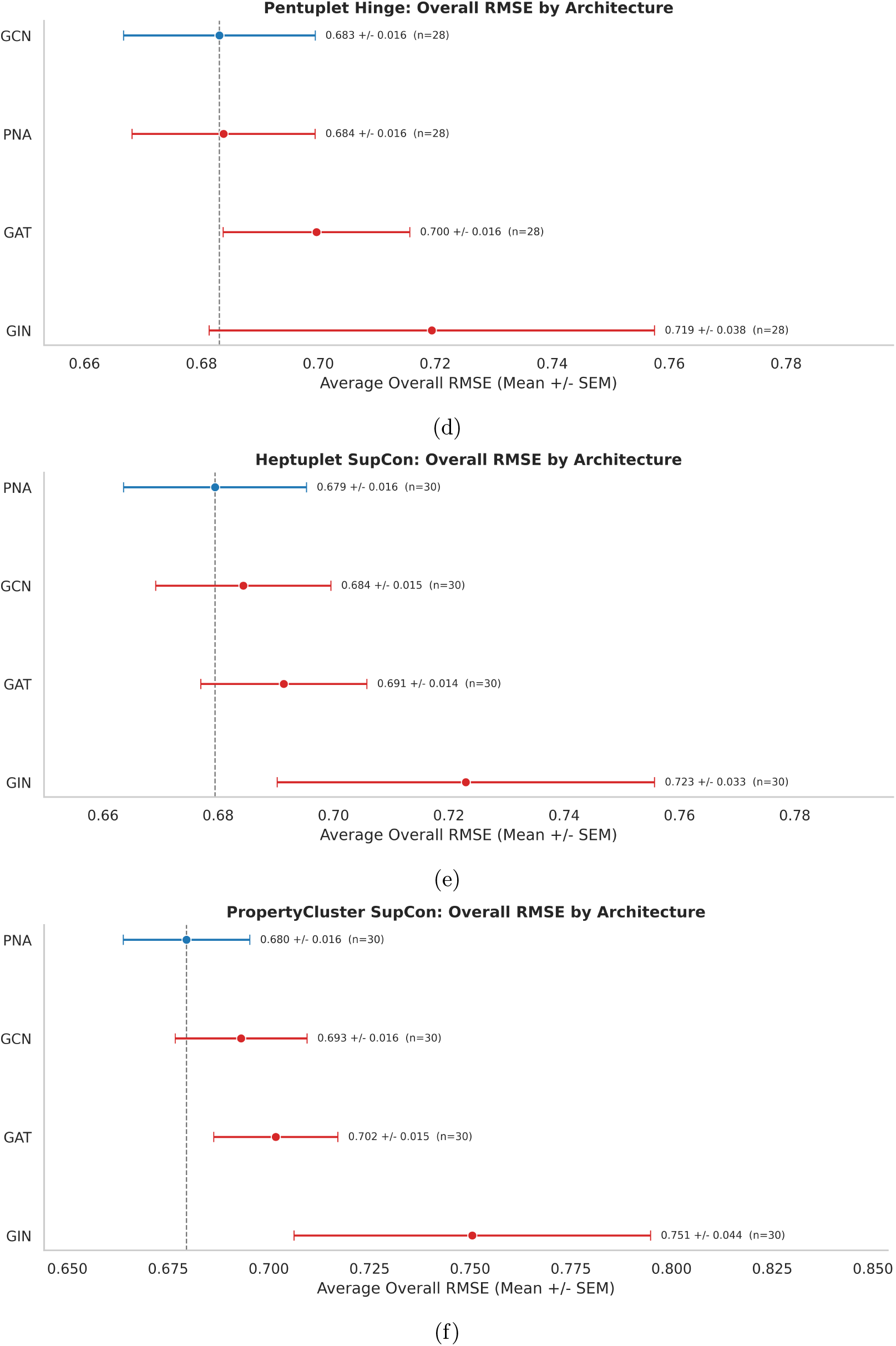
Architecture comparison for each loss function by overall test RMSE over the 30 ChEMBL activity cliff datasets.

**Figure 5:**
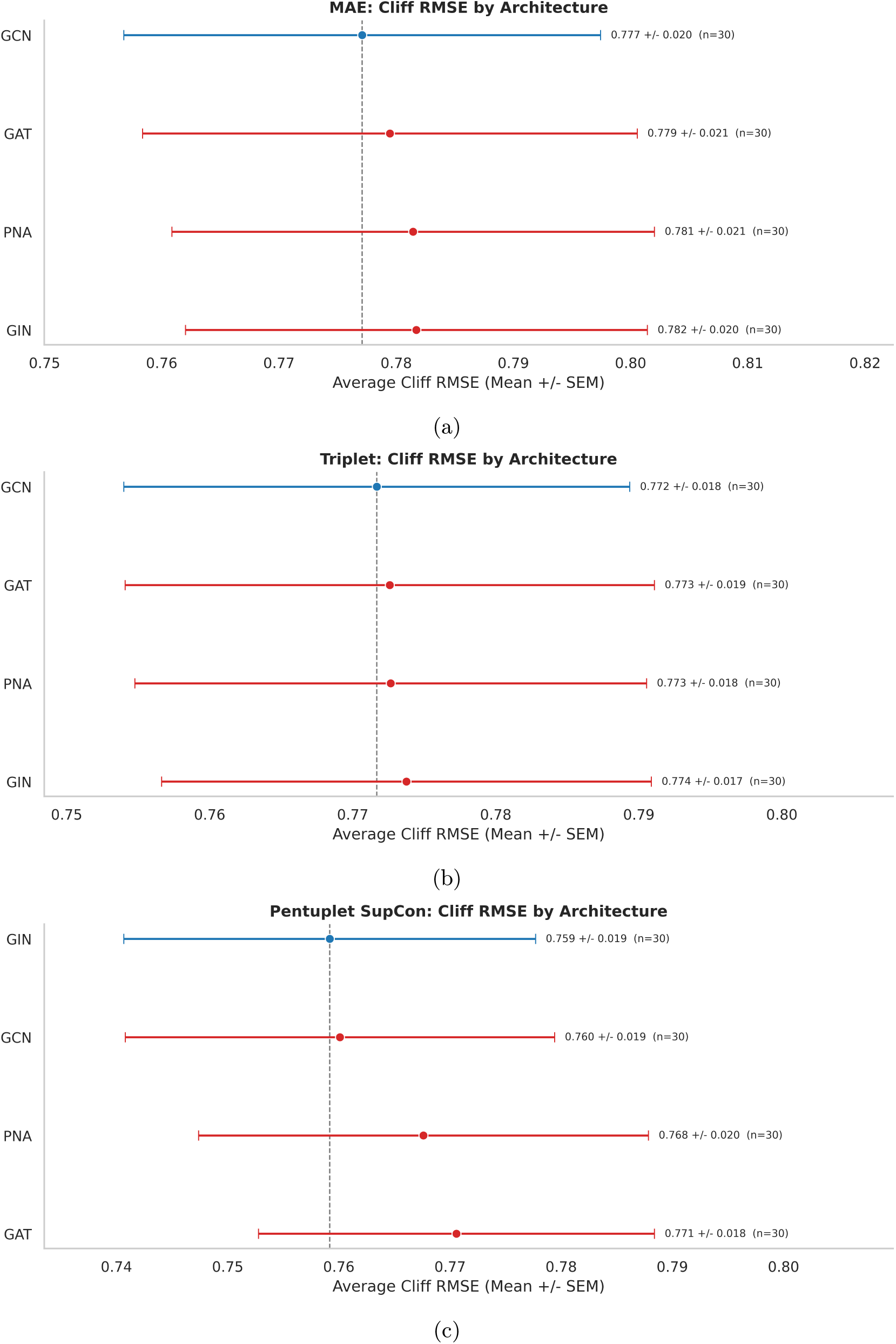
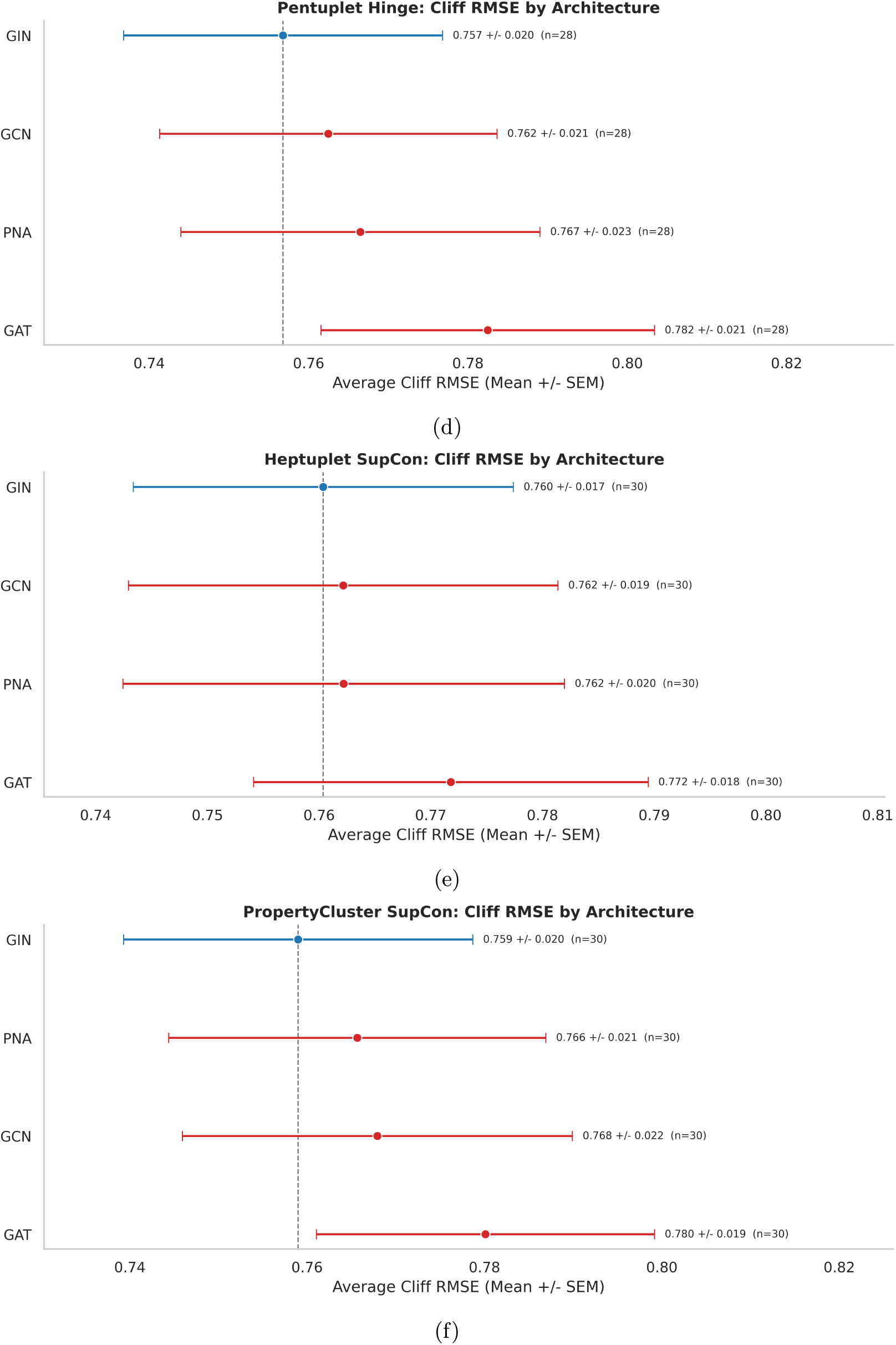
Architecture comparison for each loss function by cliff test RMSE over the 30 ChEMBL activity cliff datasets.

### Best (architecture,loss) for each dataset

We compared the best architecture and loss combination to the values reported in the literature in the ACANet benchmark. Four architectures: PNA, GIN, GCN and GAT were used and six loss functions: MAE, triplet, pentuplet SupCon, pentuplet hinge, heptuplet SupCon and propcluster SupCon were used in this study. The best combination was used to compare with the benchmarks. Figure 6 provides a comprehensive benchmarking of the best dataset specific architecture and loss combinations against the values reported in the ACANet benchmarking studies. The resulting bars which order the datasets by the best multiarch RMSE demonstrate competitive performance, even outperforming some of the previous benchmark scores, which is especially evident in the cliff test set (Figure 6(b)). Utimately, this comparison reinforces the central premise of this study that the immense structural diversity of activity cliffs necessitates highly customized, adaptive modeling strategies which goes beyond using contrastive approaches to derive meaningful structural insights that heavily drive these potency changes.

**Figure 6:**
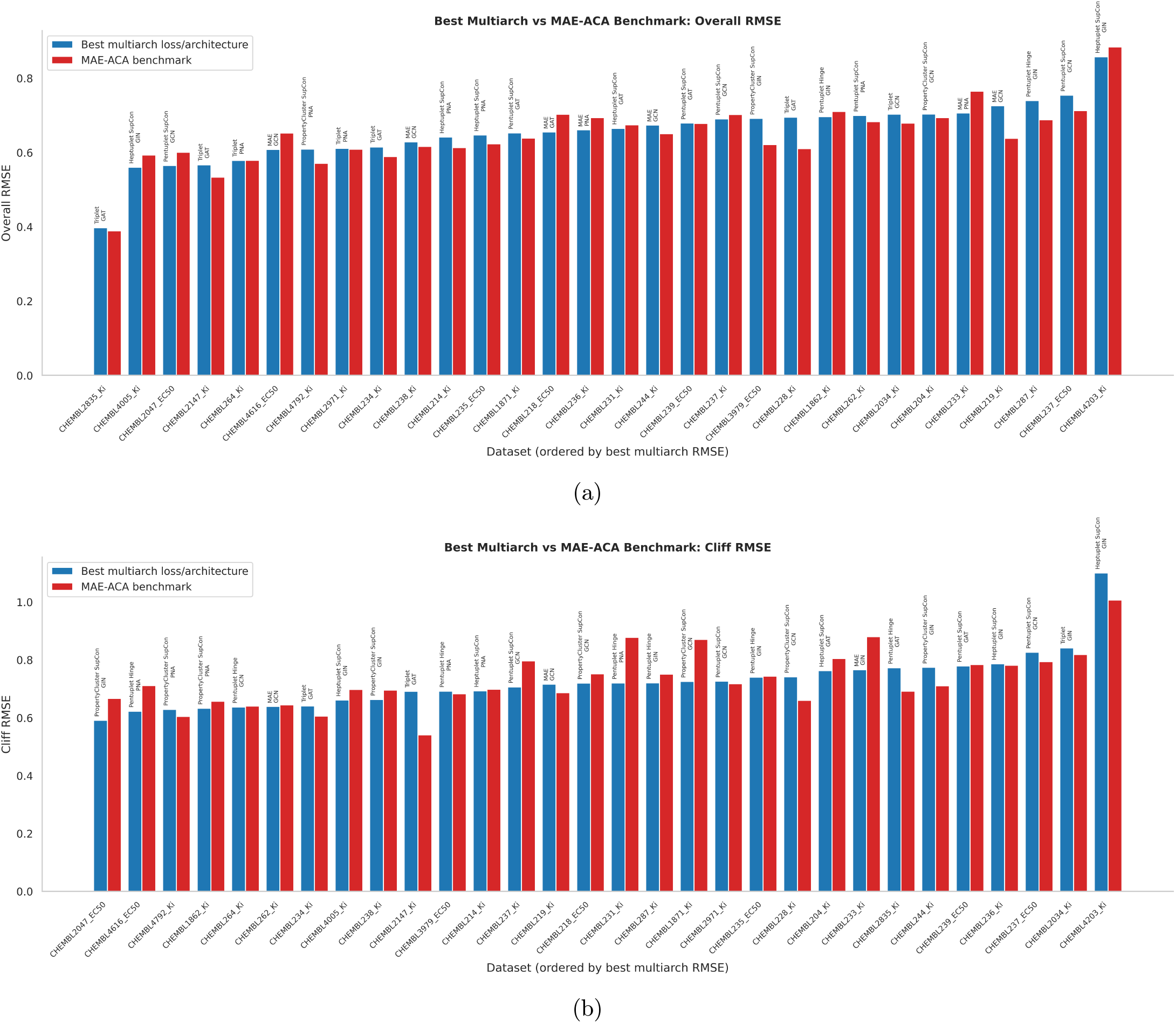
Best architecture and loss function combination to the previously reported values from the ACANet activity cliff benchmarks.

### Latent Space Embeddings

The embedding space was projected in 2D using PCA for the CHEMBL234_Ki cliff test set for the PNA architecture trained on all six loss functions. One can immediately observe qualitatively how MAE fails to distinguish by property in the latent space, while the addition of loss terms (Figure 7(b) to 7(f)) appears to make a significant difference in separating the embeddings by property value. By observing the corresponding Spearman correlations, it is observed that the projections trained on MAE performed poorly, showing no property correlation with distance in the latent space. Losses like the Triplet and Pentuplet SupCon indicated a relatively strong performance by the contrastive component playing a crucial role in separating molecules with different activities. This proves the importance of the contrastive component in the loss which helps to distinguish similar molecules. On the other hand, these results also highlight an interesting flaw in GNNs trained on contrastive learning-based losses that although they are effective in improving the interpretability of latent space embeddings by being able to project property-similar molecules closer together and prevent representation collapse, there are no clear gains when it comes to predicting the actual properties of these molecules.

**Figure 7:**
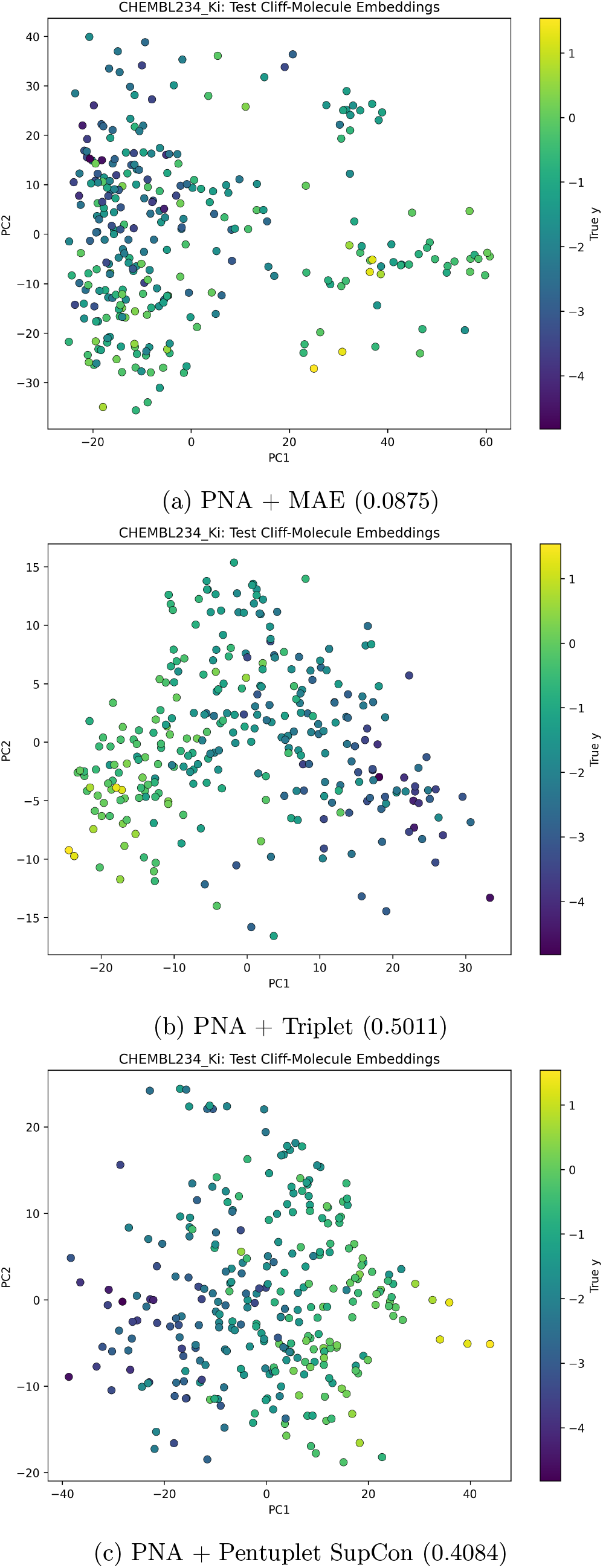
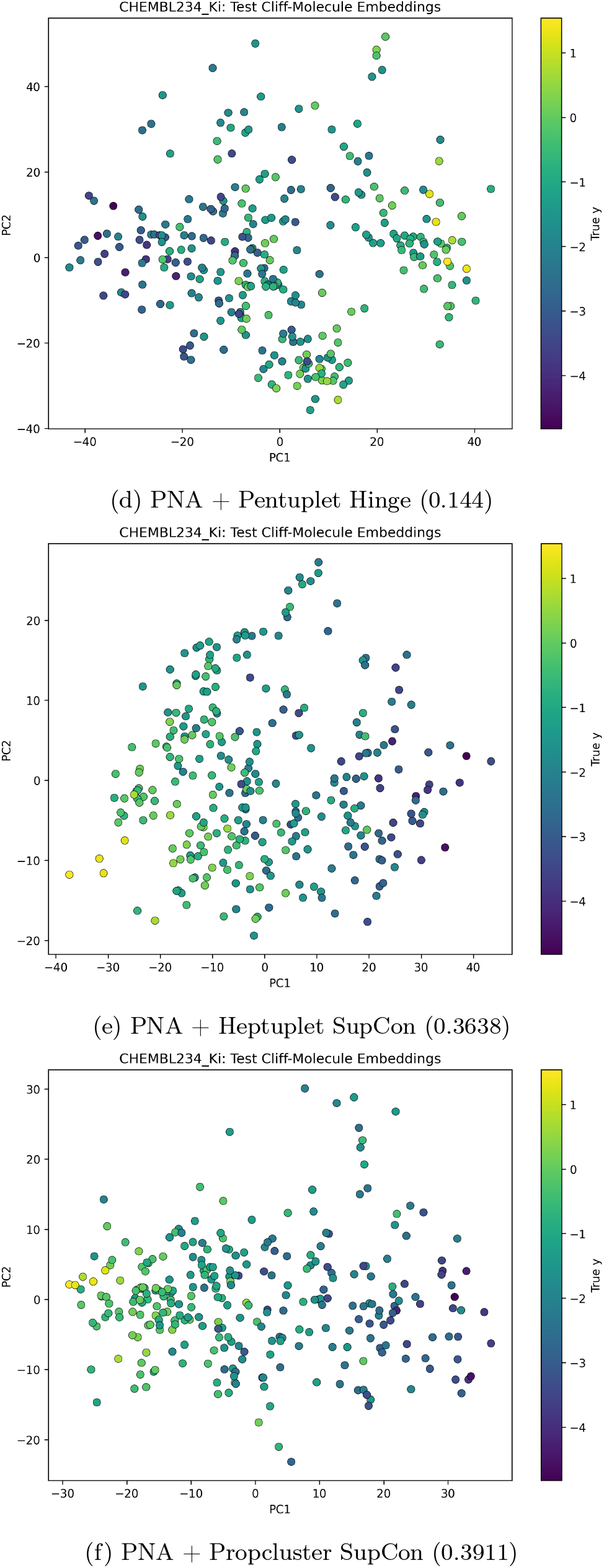
PCA projections of the embedding space of the cliff test set of CHEMBL234_ Ki along with the Spearman correlation between embedding distance and property.

## Conclusions

In this study, we evaluated the efficacy on integrating contrastive learning based loss functions into various GNN architectures to address the problem of predicting properties of activity cliff molecules. Evaluated against 30 ChEMBL subsets from the MoleculeACE benchmark repository, our results indicate that SupCon based approaches like Pentuplet, Heptuplet and PropCluster loss were effective in reducing error margins on the highly challenging cliff datasets. Furthermore, the choice of GNN backbone is important as PNA and GCN exhibited robust stability across loss functions.

The most significant contribution lies in the interpretability of the embedding space, where the PCA projections of the embeddings trained using contrastive based losses show a stronger Spearman correlation, effectively mitigating representation collapse by segregating molecules by property and generating a latent space that respects these sharp discontinuities. However, this interpretability does not directly translate to accurate property prediction, underscoring that the performance remains heavily dataset dependent. Ultimately, navigating the structure-activity landscape of activity cliffs requires not just adaptive training objectives, but highly customized training strategies that can leverage the insights derived from contrastive-learning based loss functions to guide targeted and informed lead optimization.

## Acknowledgements

We thank support from the National Institute of General Medical Sciences and the National Institutes of Health under award number R35GM150620.

